# Multi-omic analysis of photoreceptor alterations during early-onset retinal degeneration in *Mfrp^-/-^* mice

**DOI:** 10.1101/2025.08.02.668277

**Authors:** DaNae R. Woodard, Anil Chekuri, Kelsey Dang, Pooja Biswas, Swanand Koli, Justin Buchanan, Lin Lin, Sebastian Preissl, Allen Wang, Radha Ayyagari

**Affiliations:** Shiley Eye Institute, University of California San Diego, La Jolla, CA USA; Schepens Eye Research Institute, Massachusetts Eye and Ear Infirmary, Harvard Medical School, Boston, MA, USA.; Center for Epigenomics, University of California San Diego, La Jolla, CA

## Abstract

Loss-of-function mutations in the membrane frizzled-related protein (*MFRP*) gene are associated with autosomal recessive ocular disorders such as nanophthalmos, posterior microphthalmia, and retinitis pigmentosa, yet the molecular mechanisms underlying *MFRP*-related retinal degeneration remain poorly defined. To investigate these mechanisms, we generated and characterized *Mfrp^-/-^* mice and conducted an integrated multi-omic analysis to map the onset and progression of transcriptional changes during retinal degeneration. Using longitudinal *in vivo* imaging, histological and immunohistochemical analysis, qRT-PCR, single-nucleus RNA sequencing (snRNA-seq), snATAC seq (single-nucleus Assay for Transposase-Accessible Chromatin using sequencing) and spatial transcriptomics (GeoMx DSP), we identified early and progressive photoreceptor loss in *Mfrp^-/-^* mice, beginning by 28d. Retinal thinning and autofluorescent deposits in *Mfrp^-/-^* mice were accompanied by decreased expression of rod and cone opsins. Transcriptomic profiling by snRNA-seq revealed distinct, photoreceptor-specific gene expression changes, including early upregulation of inflammatory and stress-related genes and later downregulation of genes involved in chromatin remodeling, synaptic function, and neurodevelopment. Spatial transcriptomic analysis of the outer nuclear layer (ONL) revealed disruptions in mitochondrial function, mTORC1 signaling, and G protein–coupled receptor pathways. These findings define a temporally and spatially coordinated program of degeneration triggered by *Mfrp* loss and demonstrate that *Mfrp* is required for the maintenance of photoreceptor integrity and gene regulatory homeostasis during retinal development. Our work establishes *Mfrp^-/-^* mice as a robust model for dissecting photoreceptor vulnerability and provides a transcriptomic atlas of disease progression with potential relevance for therapeutic development in *MFRP*-associated retinal dystrophies.

## INTRODUCTION

Photoreceptors are highly specialized neurons responsible for initiating vision by converting light into neural signals. The structure and function of photoreceptors are critically supported by the adjacent retinal pigment epithelium (RPE), which maintains photoreceptor health through the phagocytosis of outer segments, nutrient transport, and visual cycle activity^1–3^. Disruption of the photoreceptor-RPE interface underlies inherited retinal diseases, often resulting in progressive vision loss^4^.

Mutations in membrane frizzled-related protein (*MFRP*) have been implicated in a diverse spectrum of autosomal recessive ocular disorders such as nanophthalmos, posterior microphthalmia, foveoschisis, and retinitis pigmentosa^5–10^. *MFRP* encodes a type II transmembrane protein expressed predominantly in the RPE and ciliary body, where it is thought to contribute to ocular axial length regulation and photoreceptor maintenance^11–16^. Despite these associations, the molecular function of MFRP and the mechanisms by which its loss leads to photoreceptor degeneration remain poorly understood. To investigate these mechanisms, we generated and characterized a novel homozygous *Mfrp* knockout mouse model (*Mfrp^-/-^*). *Mfrp^-/-^*mice recapitulate key features of human MFRP-associated retinal degeneration, including early-onset photoreceptor loss, retinal thinning, and fundus abnormalities. While previous studies have described retinal degeneration in *Mfrp* mutant mice^12,16–18^, a comprehensive single-cell transcriptomic analysis of disease progression has not been performed.

Herein, we present an integrated multi-omic analysis of the *Mfrp^-/-^* retina to define the cellular and molecular landscape of early-onset retinal degeneration. Single-nucleus RNA sequencing (snRNA-seq) was used to profile transcriptomic changes across retinal cell types, with a focus on photoreceptors. These findings were spatially resolved using GeoMx Digital Spatial Profiling, enabling high-resolution mapping of gene expression changes within the photoreceptor outer nuclear layer (ONL). In parallel, ATAC-seq was performed to assess chromatin accessibility and uncover regulatory changes associated with photoreceptor dysfunction. Together, these approaches revealed early transcriptional and epigenetic disruptions in both rods and cones, including dysregulation of genes involved in phototransduction, metabolic homeostasis, and cellular stress pathways. Our findings demonstrate that *Mfrp* deficiency drives early, cell type– specific degeneration and redefines the transcriptional landscape of the degenerating retina. This work provides a molecular framework for understanding MFRP-associated retinopathy and a foundation for future therapeutic discovery.

## RESULTS

### Mfrp^-/-^ mice exhibit progressive retinal degeneration characterized by fundus abnormalities, and retinal thinning, and photoreceptor loss

To evaluate the impact of *Mfrp* deletion on retinal structure and integrity, a comprehensive analysis of the fundus, retinal thickness, and histological organization was performed on *Mfrp^-/-^*and Wt mice (Fig 1). Fundus autofluorescence imaging revealed retinal changes in *Mfrp^-/-^*mice that were not observed in Wt controls. In Wt mice, no fundus abnormalities were detected at 1-2mo, 2.5-3mo, and 5-6mo (Fig 1A, top row). In contrast, *Mfrp^-/-^* mice displayed a progressive increase in autofluorescent spots throughout the retina at 1-2mo of age, which became increasingly prominent by 5-6mo (Fig 1A, bottom row). To assess and quantify structural changes in *Mfrp^-/-^* mice, optical coherence tomography (OCT), was performed in 2.5mo *Mfrp^-/-^* mice and compared to 2.5mo Wt controls (Fig 1B). Representative images from the central retina revealed a substantially thinner retinal architecture in 2.5mo *Mfrp^-/-^* mice compared to 2.5mo Wt mice (Fig 1B). Quantification of the total retinal thickness in 2.5mo *Mfrp^-/-^* mice confirmed a significant decrease in both the central retina (*Mfrp^-/-^*= 152 ± 4.7 μm vs. Wt = 231.2 ± 8.5 μm; ****p < 0.0001) and peripheral retina (*Mfrp^-/-^* = 159 ± 3.9μm vs. Wt = 237.5 ± 3.6 μm ****p < 0.0001) (Fig. B, bottom).

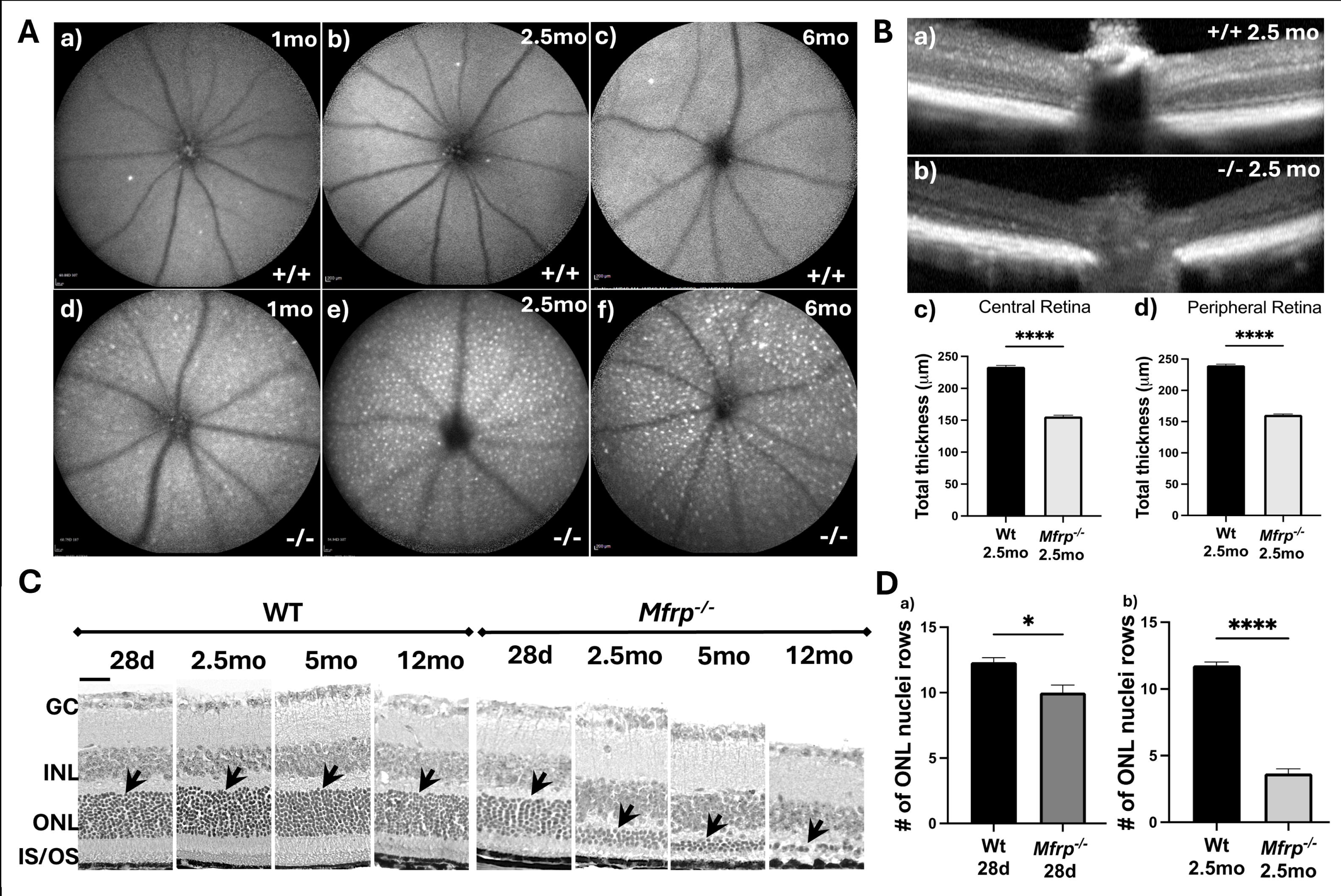

To further investigate the nature of retinal thinning in *Mfrp^-/-^*mice, histological retinal cross-sections from *Mfrp^-/-^* and Wt mice at postnatal day 28 (28d), 2.5mo, 5mo, and 12mo of age (Fig 1C). Wt mice displayed a preserved retinal structure at each timepoint with distinguishable layers including the ganglion cell layer (GCL), inner nuclear layer (INL), outer nuclear layer (ONL), and inner and outer segments (IS/OS) of photoreceptors. In contrast, *Mfrp^-/-^* mice showed significant and progressive degeneration of the photoreceptor ONL (Fig 1C). At 28d, *Mfrp^-/-^* mice displayed a modest reduction in ONL thickness (Fig 1C). By 2.5mo, the ONL was severely diminished, and by 12mo, only 1 row of nuclei was detected, along with the disruption of the IS/OS region in *Mfrp^-/-^* mice (Fig 1C). Quantitative assessment of ONL thickness in *Mfrp^-/-^* mice confirmed fewer rows of photoreceptor nuclei at 28d (*Mfrp^-/-^* = 10 ± 2 vs. Wt = 12.3 ± 1.1 vs. *p < 0.05) (Fig 1D, left panel) and dramatic reduction in photoreceptor nuclei at 2.5mo (*Mfrp^-/-^* = 3.6 ± 1.1 vs. Wt = 11.7 ± 0.7 vs. ****p < 0.0001) (Fig 1D, right panel). Together, these findings demonstrate that complete loss of *Mfrp* in the retina results in early and progressive retinal degeneration.

### Loss of Mfrp alters photoreceptor opsin expression

Given the observed ONL thinning in *Mfrp^-/-^* mice, investigation of photoreceptor morphology and opsin expression was assessed via immunohistochemistry and qRT-PCR in *Mfrp^-/-^* and Wt mice at 28d and 2.5mo (Fig 2). Immunofluorescent labeling for photoreceptor-specific markers (Rhodopsin, M-opsin, and S-opsin) revealed distinct differences between *Mfrp^-/-^*and Wt retinas (Fig 2A). At 28d, Wt mice showed robust RHO immunoreactivity in the OS (Fig 2A, top left panel). In contrast, *Mfrp^-/-^*mice displayed markedly reduced RHO in the OS (Fig 2A, top second panel). This phenotype was further exacerbated at 2.5mo, with *Mfrp^-/-^* mice showing minimal RHO expression compared to 2.5mo Wt controls (Fig 2A, top right panels). Similarly, both M-opsin and S-opsin expressing cones revealed significant disruptions in the OS structure. At 28d, M-opsin and S-opsin expressing cone OS in *Mfrp^-/-^* mice were similar in morphology to 28d Wt mice. However, by 2.5mo, *Mfrp^-/-^* mice showed a drastic shortening of M-opsin and S-opsin expressing cones, which resembled small puncta (Fig 2A, middle and bottom panels).

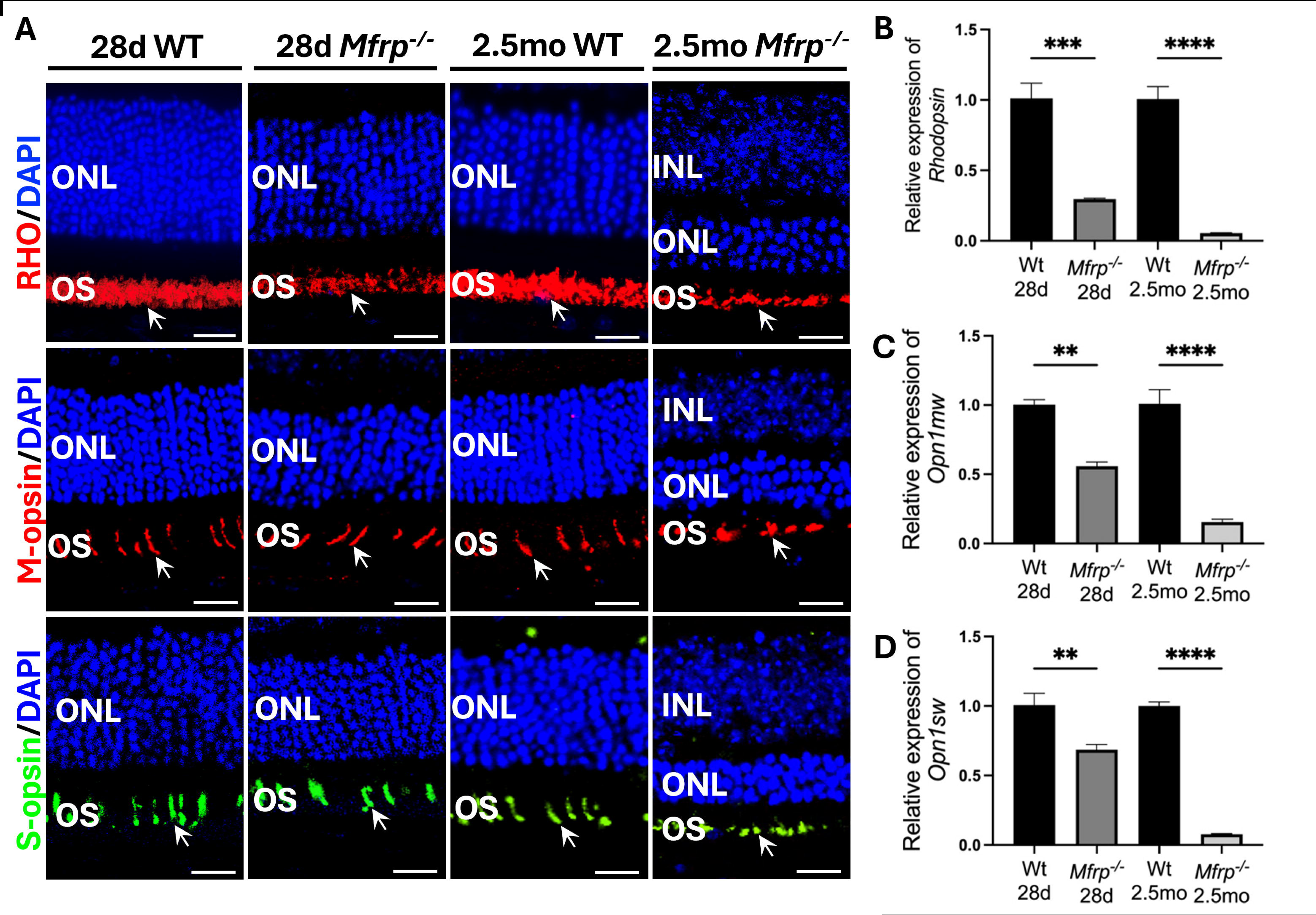

Analysis of photoreceptor-specific transcripts corroborated the immunohistochemistry findings (Fig 2B-D). Expression of rod-specific *Rho* gene expression was significantly decreased in *Mfrp^-/-^* mice at 28d and 2.5mo compared to age-matched Wt mice (Fig 2B). Cone-specific opsins, *Opn1mw* and *Opn1sw*, were significantly reduced in *Mfrp^-/-^* mice compared to age-matched Wt mice at 28d and 2.5mo (Fig 2B, C). *Opn1mw* expression was reduced by ∼50% at 28d and ∼80% in *Mfrp^-/-^* mice compared to Wt controls (Fig 2B). A similar trend was observed for *Opn1sw* (Fig 2C). Together, these findings demonstrate that *Mfrp* deficiency leads to progressive degeneration of rod and cone OS and a marked downregulation of photoreceptor-specific genes, confirming a critical role of *Mfrp* in photoreceptor maintenance.

### Single-nucleus RNA sequencing reveals distinct retinal cell populations in Mfrp^-/-^ mice

To investigate how *Mfrp* loss affects retinal cell populations and gene expression at the single cell level, single-nucleus RNA sequencing (snRNA-seq) was performed on retinas dissected from Wt and *Mfrp^-/-^*mice at 28d and 2.5mo (Fig 3). Following tissue dissection, nuclei were extracted and barcoded for Illumina sequencing (Fig 3A). Dimensionality reduction and clustering were performed using Uniform Manifold Approximation and Projection (UMAP) to define cell type identities (Fig 3A). Clustering analysis of 57,420 nuclei from Wt and *Mfrp^-/-^* retinas identified 19 diverse cell types (Fig 3B). Distinct clusters were annotated based on canonical marker gene expression that included photoreceptors (rods and cones), bipolar cells (ON- and OFF-cone bipolar, rod bipolar) amacrine cells (GABAergic, glycinergic, and All-amacrine), Müller glia, retinal ganglion cells (RGCs), horizontal cells, microglia, astrocytes, fibroblasts, endothelial cells, melanocytes, smooth muscle cells (SMCs), pericytes, and retinal pigment epithelium (RPE) (Fig 3B).

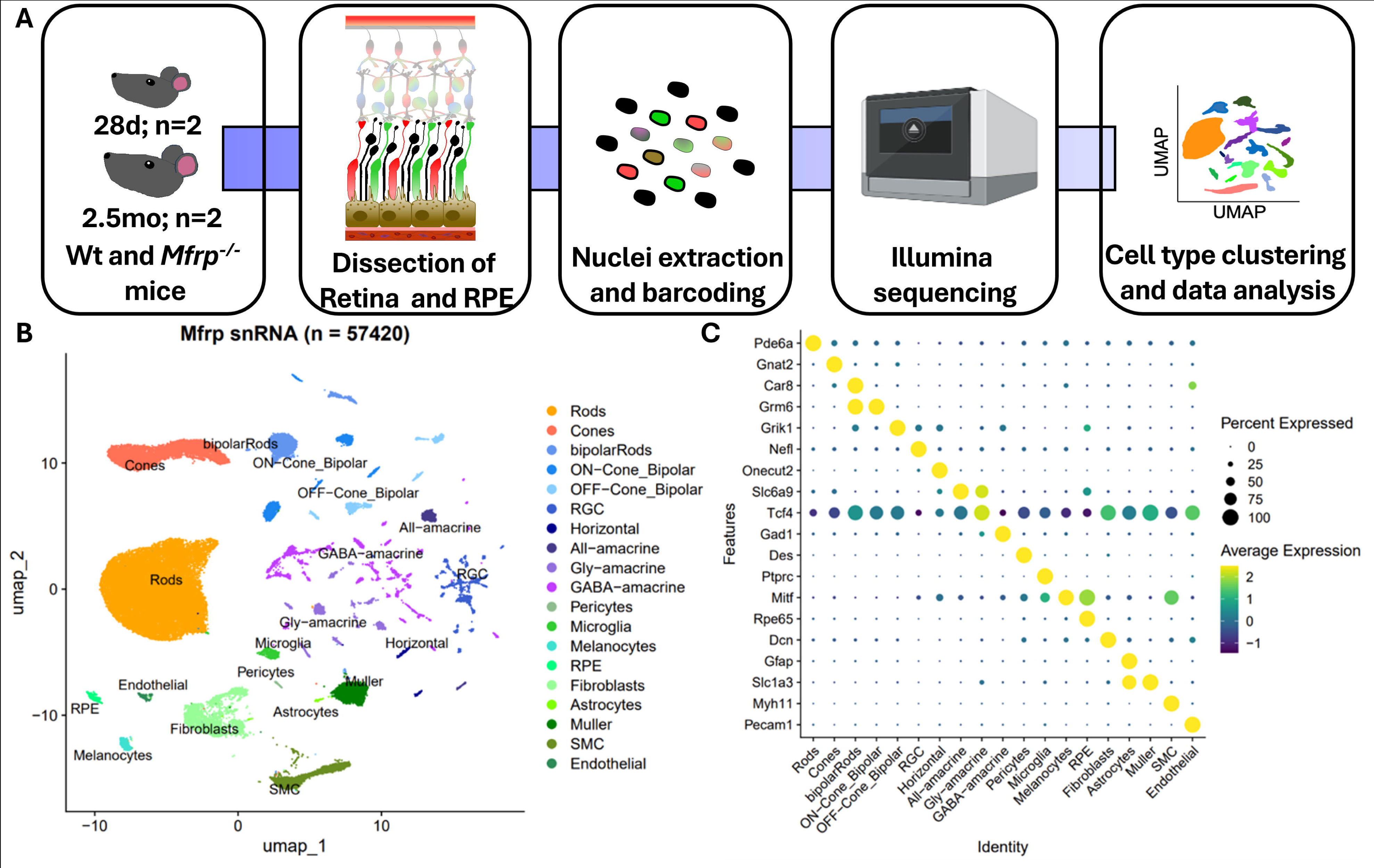

To confirm cell type identities, expression of well-characterized marker genes were examined across the clusters (Fig 3C). Expression of canonical photoreceptor genes validated the identity of rod and cone clusters (*Pde6a* and *Gnat2*, respectively) (Fig 3C). Bipolar cell populations expressed *Grm6*, *Grik1*, and *Onecut2*, while amacrine subtypes were delineated by *Gad1* and *Slc6a9* (GABAergic and glycinergic, respectively) (Fig 3C). *Rpe65* and *Mitf* expression was restricted to RPE cells, while Müller glia, microglia, endothelial cells, and other non-neuronal cell types displayed expression of canonical markers including *Slc1a3*, *Pecam1*, *Ptprc*, and *Dcn* (Fig 3C).

### Differential gene expression of rod and cone photoreceptors in Mfrp^-/-^ mice

To specifically identify transcriptomic changes in rod and cone photoreceptors due to the loss of *Mfrp*, differential gene expression analysis was performed to compare Wt and *Mfrp^-/-^* mice at 28d and 2.5mo of age. Volcano plots were generated to visualize significant gene expression changes (adjusted p <0.05, log2 fold change >1) (Fig 4). At 28d, *Mfrp^-/-^* rod photoreceptors exhibited significant gene expression changes relative to Wt mice (Fig 4A).

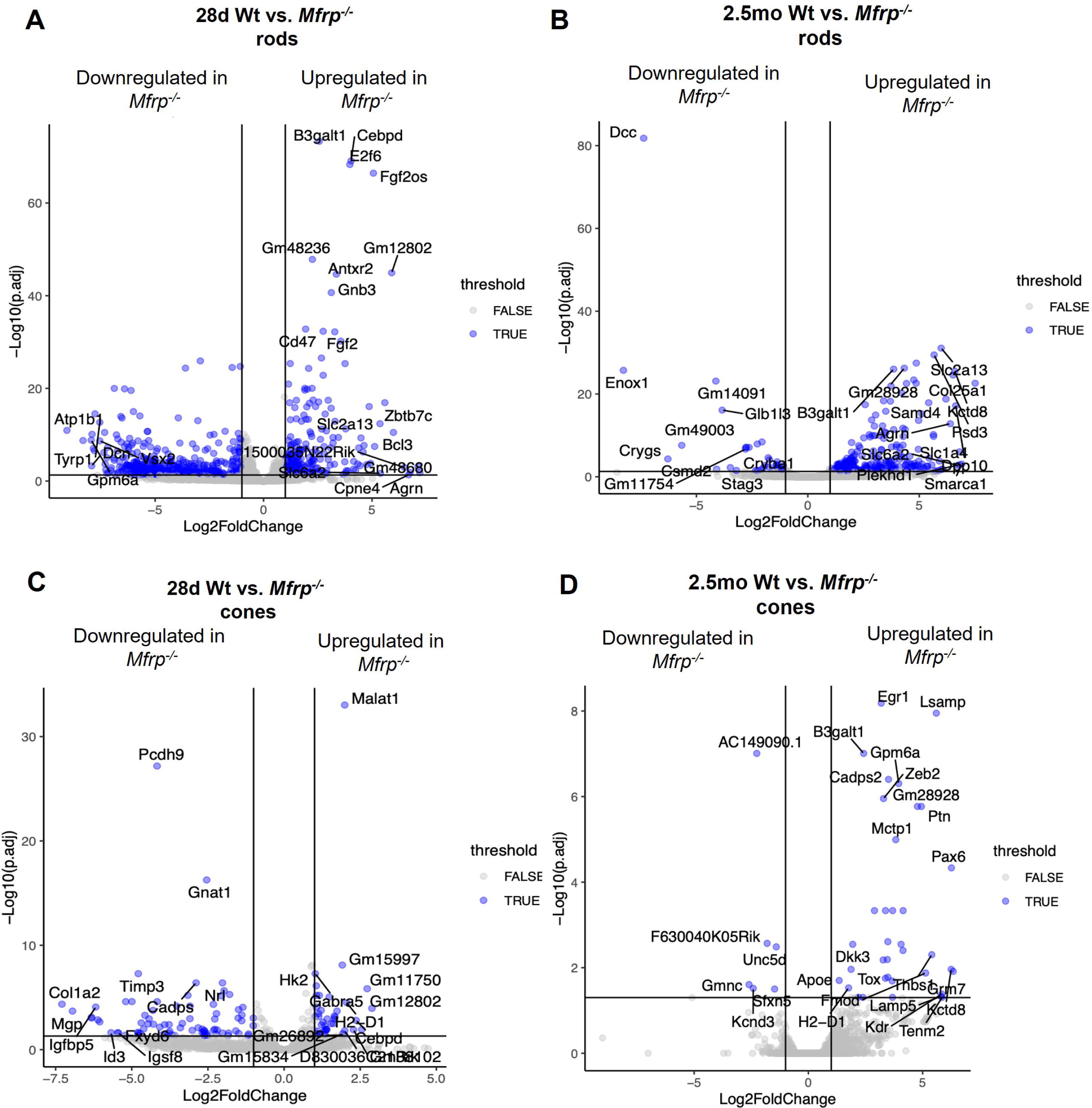

Upregulated genes in 28d *Mfrp^-/-^* rods were enriched in processes related to cytokine signaling and stress response (Fig 4A). Notable upregulated genes include *Cebpd*, transcription factor associated with inflammatory response and reactive gliosis^19^; *B3galt1*, involved in glycosylation and cellular adhesion^20^; *Fgf2os*, a member of the fibroblast growth factor family with roles in neurodevelopment^21^; and *Gnb3*, a G-protein subunit implicated in visual signal transduction^22^ (Fig 4A). Conversely, downregulated genes in 28d *Mfrp^-/-^* rods were primarily associated with photoreceptor metabolism and homeostasis such as *Atp1b1* and *Gpmls4* (Fig 4A)*. Atp1b1* which encodes the β-subunit of Na^+^/K^+^ ATPase and plays a critical role in ion transport essential for photoreceptor excitability^23,24^. *Gpmls4* is a mitochondrial gene related to oxidative phosphorylation^25^. These early transcriptional changes suggest that absence of *Mfrp* impacts rod photoreceptor homeostasis during the developmental stage.

At 2.5mo, the transcriptional landscape in in *Mfrp^-/-^*rods shifted towards downregulation, particularly in genes associated with chromatin organization and cellular architecture (Fig 4B). Key downregulated genes include *Smarca1*, a chromatin remodeling factor essential for gene retinal development and photoreceptor maintenance^26^; *Stag3*, involved in cohesion-mediated chromatin structure and genome integrity^27^; and *Gm28928*, a non-coding RNA with potential regulatory function (Fig 4B). In contrast, upregulated genes such as *Dcc*, a netrin receptor involved in apoptosis and axon guidance^28,29^, and *Enox1* (linked to redox balance^30^), may suggest ongoing cellular stress or compensatory signaling (Fig 4B). These findings are indicative of progressive rod photoreceptor degeneration and cellular reprogramming in 28d and 2.5mo *Mfrp^-/-^* mice.

In *Mfrp^-/-^* cone photoreceptors, transcriptional alterations were already apparent at 28d (Fig 4C). Downregulated genes in *Mfrp^-/-^*cones included *Pcdh9* (encodes for a collagen-associated protein that affects cell adhesion^31^); *Gnat1*, a rod transducing subunit often misregulated in retinal disease^32^; *Timp3*, which modules extracellular matrix turnover and implicated in Sorsby fundus dystrophy^33,34^; and *Col1a2*, a collagen subunit critical for structural integrity^35^ (Fig 4C). Upregulated genes in *Mfrp^-/-^*cones included *Malat1*, a long non-coding RNA frequently associated with oxidative stress responses in retina cells^36^; uncharacterized long non-coding RNAs such as *Gm12802* and *Gm15997*; and *Hk2,* an essential enzyme in aerobic glycolysis^37^(Fig 4C).

*Mfrp^-/-^* cones at 2.5mo exhibited a broader and more severe pattern of dysregulation, with significant downregulation of genes involved in neurodevelopment, synaptic organization, and immune regulation. Key downregulated genes included *Egr1*, an immediate early gene important for synaptic plasticity and retinal development^38^; *Gpm6a*, which is involved in neurite outgrowth and plasticity^39^; *Pax6*, a master transcription factor in eye development and retinal identity^40,41^; and *Ptn*, a neurotrophic factor supporting neuron survival^42^. Upregulated genes included *FG630040K05Rik*, a long non-coding RNA; *Unc5d*, a netrin receptor associated with axon repulsion and apoptosis^43,44^; and *Dkk3*, a Wnt signaling modulator implicated in retinal vascular and neurodegenerative disease^45^. Together, these findings indicate that *Mfrp* deficiency leads to progressive and divergent patterns of gene dysregulation in rod and cone photoreceptors at 28d and 2.5mo.

### Transcriptomic pathway analysis reveals dysregulation of key biological processes in Mfrp^-/-^ photoreceptors

To further investigate the biological processes altered by *Mfrp* deficiency, gene set enrichment analysis (GSEA) was performed on differentially expressed genes (DEGs) identified in rods and cones from 28-day-old and 2.5-month-old mice using DESeq2. DEGs with an adjusted *p*-value < 0.05 and absolute log_₂_ fold-change ≥ 1 were ranked by log_₂_ fold-change and analyzed using the clusterProfiler R package (v4.0). Heatmaps revealed distinct sets of significantly upregulated and downregulated pathways at 28d and 2.5mo (Fig 5). In 28d *Mfrp^-/-^* rods, significantly upregulated pathways included regulation of miRNA transcription, interleukin-10 production, and response to ischemia (Fig 5A), suggesting an early activation of inflammatory and stress-response mechanisms. At 2.5mo, *Mfrp^-/-^* rods showed significant enrichment of upregulated pathways that include visual system development, synapse organization, and response to fatty acid (Fig 5A), suggesting an occurrence of plasticity and metabolic adaptation. In 28d *Mfrp^-/-^* cones, significantly enriched upregulated pathways include acylglycerol metabolic process, mechanoreceptor differentiation, and phosphatidic acid metabolic process (Fig 5A), indicating early metabolic and structural adaptations in cones. By 2.5mo, *Mfrp^-/-^*cones showed enrichment of eye photoreceptor cell development, regulation of neurogenesis, and regulation of response to wound healing pathways (Fig 5A), suggesting that cones may be activating developmental and adaptive processes at this age point.

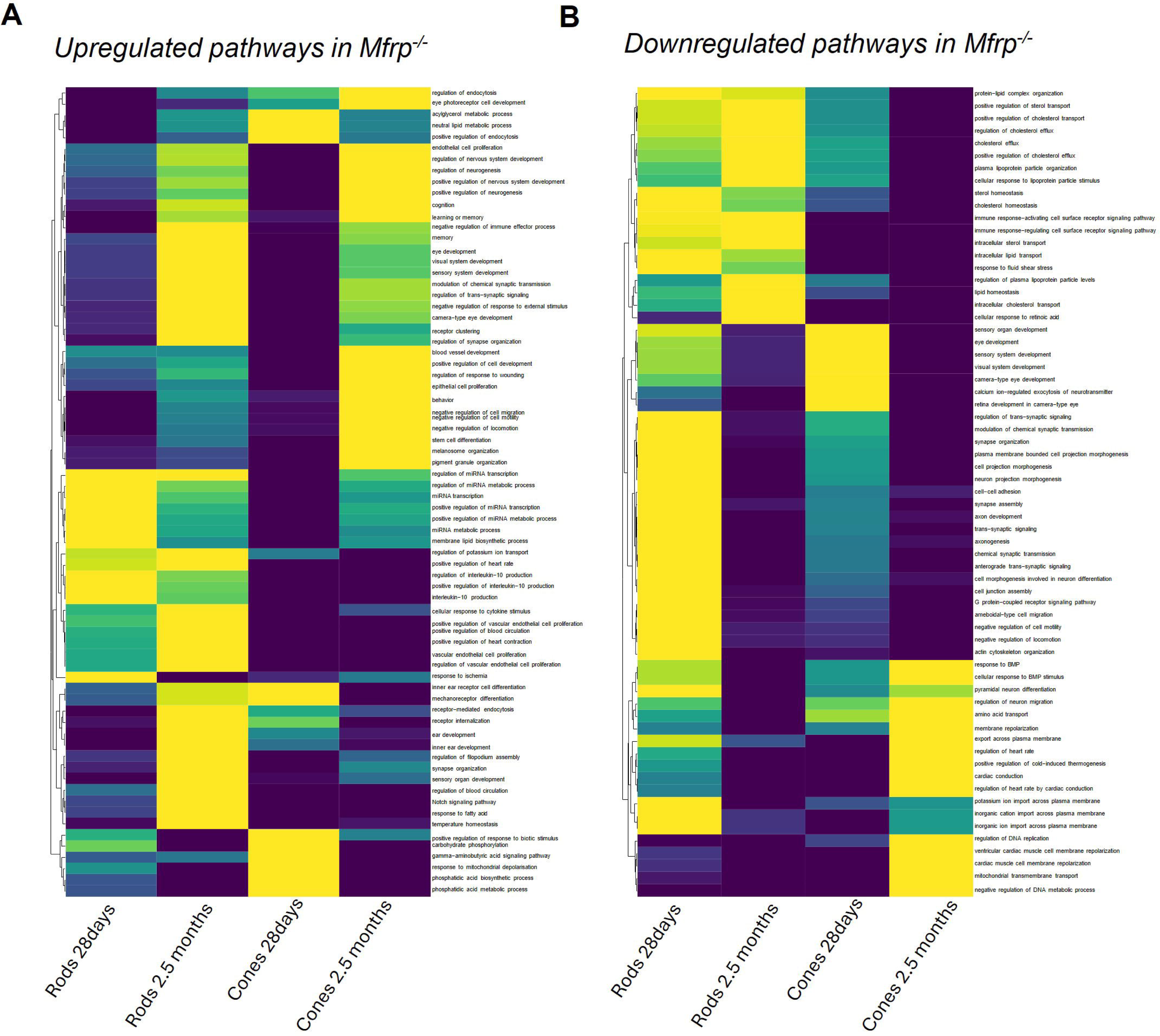

Conversely, significantly downregulated pathways in 28d *Mfrp^-/-^*rods include pathways such as cholesterol homeostasis, immune response−activating cell surface receptor signaling pathway, and chemical synaptic transmission (Fig 5B). By 2.5mo, the immune response−activating cell surface receptor signaling pathway was still detected to be downregulated as well as intracellular sterol transport in *Mfrp^-/-^* rods (Fig 5B). In 28d *Mfrp^-/-^* cones, visual system development pathways were significantly downregulated along with calcium ion−regulated exocytosis of neurotransmitter (Fig 5B), indicating an early dysregulation of cones. By 2.5mo, significantly downregulated pathways include response to BMP, amino acid transport, and mitochondrial transmembrane transport in *Mfrp^-/-^* cones (Fig 5B), suggesting the loss of key survival and metabolic processes in cones. Together, these findings highlight the dynamic and photoreceptor-specific transcriptomic alterations in the *Mfrp^-/-^*retina.

### Altered chromatin accessibility in Mfrp^-/-^photoreceptors

To explore the epigenetic mechanisms underlying photoreceptor dysfunction in *Mfrp^-/-^* mice, single cell combinatorial indexing ATAC-seq (sciATAC-seq) was performed on 28d and 2.5mo Wt and *Mfrp^-/-^* retinas. This method allows for profiling chromatin accessibility at single-nucleus resolution, offering insight in to cell-type specific regulatory changes^46^. UMAP projection of the sciATAC-seq data revealed distinct clustering of major retinal cell types, including rods, cones, Müller glia, bipolar, amacrine and ganglion cells (Fig 6A). Cell-type identity was confirmed by the expression of canonical marker genes (Fig 6B), validating the robustness of this dataset and enabling confident downstream differential accessibility analysis.

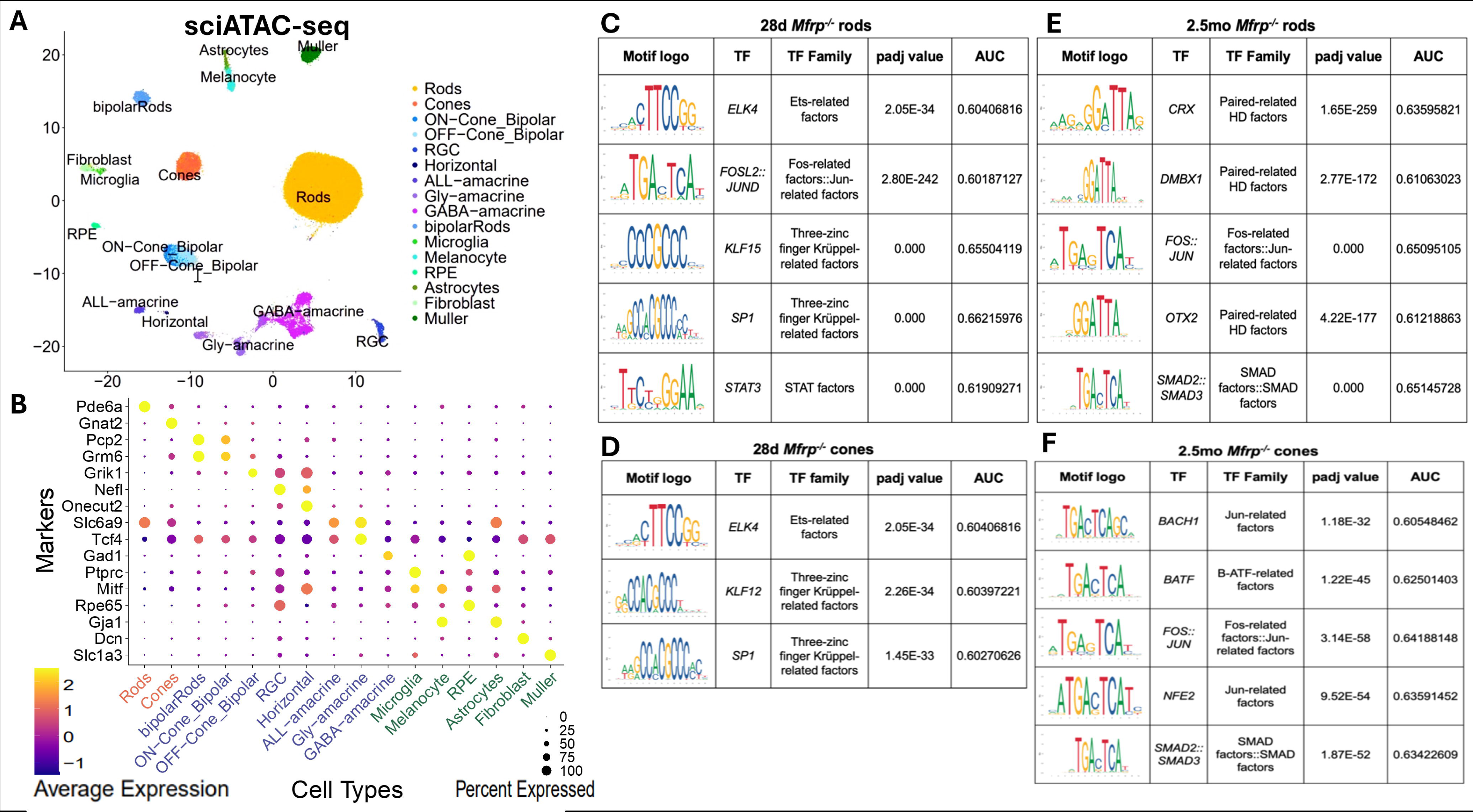

In 28d *Mfrp^-/-^* rods, widespread changes in chromatin accessibility were observed relative to age-matched Wt controls. Motif enrichment analysis of differentially accessible regions (DARs) revealed significant accessibility of binding motifs for ELK4, FOSL2::JUND (AP-1 complex), KLF15, SP1, and STAT3 (Fig 6C). ELK4, an ETS-domain transcription factor (TF), operates downstream of the ERK/MAPK pathway and has been implicated in stress-responsive gene activation^47,48^. FOSL2::JUND, representing the AP-1 complex, regulates transcription in response to oxidative stress, inflammation, and apoptosis-hallmarks of early retinal degeneration^49^. KLF15 is known for its role in regulating cellular metabolism and redox balance and is induced in response to metabolic and oxidative stress^50,51^. SP1, a broadly expressed TF involved in chromatin remodeling and DNA repair^52^. STAT3, a key effector of inflammatory signaling, is strongly associated with reactive gliosis and retinal injury^53^. In 28d *Mfrp^-/-^* cones, a similar pattern of early epigenomic alterations was observed. Motif enrichments revealed significant binding accessibility at binding motifs for ELK4, KLF12, and SP1 (Fig 6D). The repeated enrichment of these motifs suggests a shared stress-induced transcriptional state and chromatin remodeling response in 28d *Mfrp^-/-^* rods and cones.

By 2.5mo, *Mfrp^-/-^* rods exhibited extensive chromatin remodeling. Analysis of motif enrichment revealed strong enrichment of CRX, DMBX1, FOS::JUN, OTX2, and SMAD::SMAD3 binding sites (Fig 6E). While CRX and OTX2 are essential photoreceptor-specific TFs that regulate the expression of genes required for OS formation, phototransduction, and terminal differentiation^54,55^, DMBX1 is known to influence retinal patterning during development^56^. Enrichment of FOS::JUN likely suggests steady activation of stress-responses as observed at 28d *Mfrp^-/-^* rods. Enrichment of SMAD::SMAD3 indicates engagement of TGF-β signaling, a pathway implicated in neuroinflammation, fibrosis, and glial reactivity^57^. In 2.5mo *Mfrp^-/-^* cones, chromatin accessibility changes were similarly prominent. Enriched TF motifs included BACH1, BATF, FOS::JUN, NFE2, and SMAD::SMAD3 (Fig 6F). BACH1 and NFE2 are involved in oxidative stress regulation and antioxidant response^58,59^, while BATF is implicated in immune signaling^60^. Similar to 2.5mo *Mfrp^-/-^* rods, FOS::JUN and SMAD::SMAD3 was also enriched in 2.5mo *Mfrp^-/-^* cones, highlighting shared alterations in chromatin.

Collectively, these data show that loss of *Mfrp* induces profound and age-progressive changes in chromatin accessibility in photoreceptors. The enrichment of stress- and inflammation-associated TF motifs in rods and cones suggests that dysregulation in *Mfrp^-/-^* retinas is driven in part by epigenomic remodeling. These findings underscore a critical role for Mfrp in preserving the regulatory stability of photoreceptors and provide insight into early molecular events during degeneration.

### Spatial transcriptomic analysis identifies ONL-specific molecular changes in Mfrp^-/-^ mice

To dissect the spatial context of transcriptional changes, GeoMx Digital Spatial Profiling (DSP) was performed on retinal sections from Wt and *Mfrp^-/-^* mice at 28d and 2.5mo (Fig. 7A). Immunofluorescent labeling for GFAP, IBA1, and laminin revealed abnormal retinal architecture and reactive gliosis in *Mfrp^-/-^* mice, indicative of neuroinflammation and retinal stress (Fig. 7B). SYTO13 nuclear staining facilitated selection of regions of interest (ROIs) specifically within the ONL for spatial transcriptomic analysis (Fig. 7C). Pathway enrichment analysis revealed distinct alterations in biological processes, including metabolism, signal transduction, and immune system (Fig 6D), highlighting major changes in *Mfrp^-/-^* ONL.

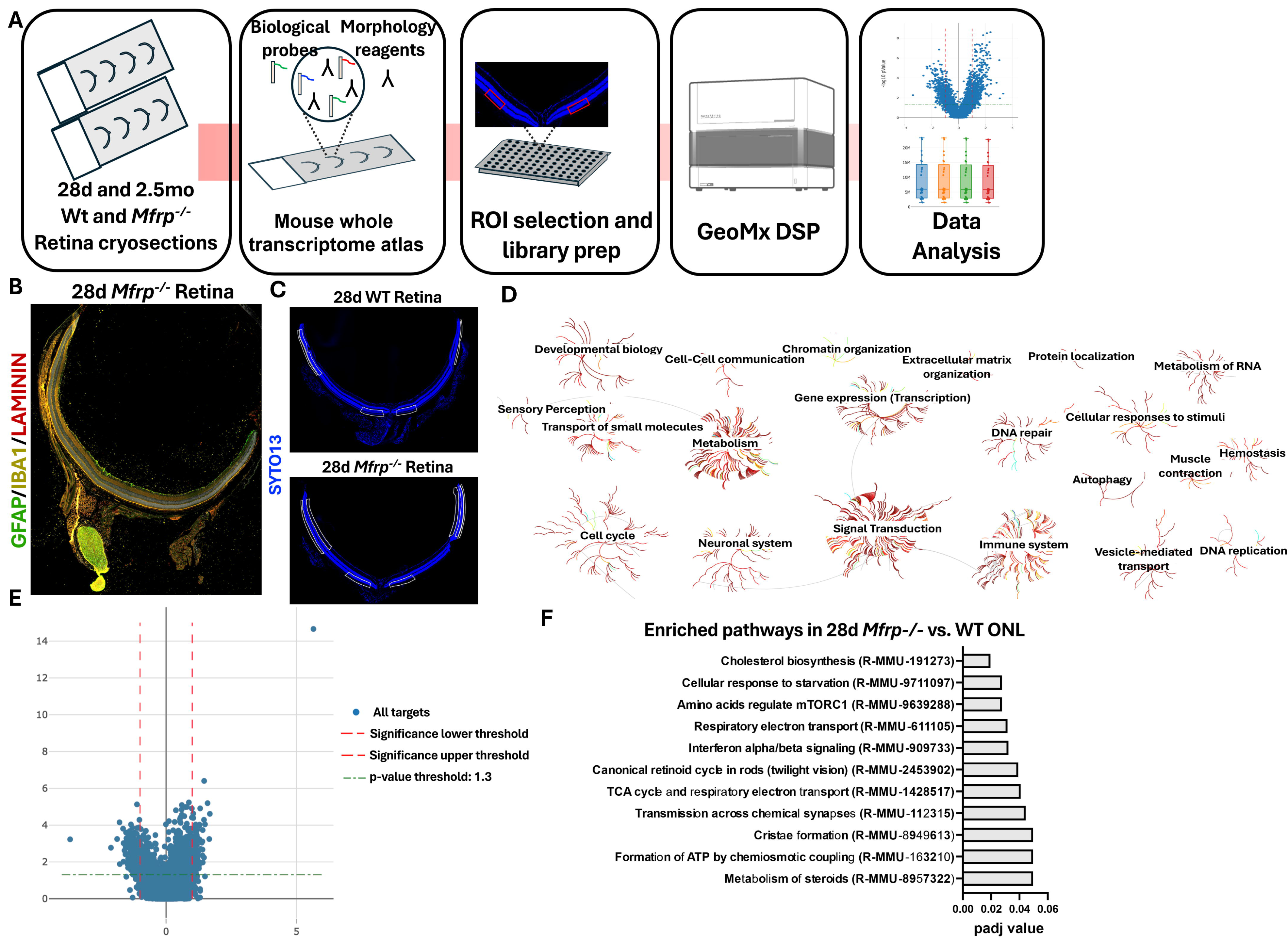

Differential expression analysis further confirmed robust transcriptional alterations in the *Mfrp^-/-^* ONL, with a subset of genes exceeding the adjusted *p*-value threshold (Fig. 7E). Among the most significantly upregulated genes was *Flp recombinase* (log_2_FC = 5.65, p = 2.19 x 10^-15^), which likely reflects the transgenic element used to genetically engineer the *Mfrp^-/-^*mice (Fig 7E). Excluding this, several biologically relevant genes were identified. *Gnb3* (log_2_FC = 1.46, p = 3.96 x 10^-7^) and *Gnb2* (log_2_FC = 1.31, p = 1.23 x 10^-5^), encoding the β subunits of heterotrimeric G protein-coupled receptor (GPCR)^61,62^, were significantly upregulated in the *Mfrp^-/-^* ONL (Fig 7E), suggesting a potential alteration in signal transduction cascades. Gpr162 (log_2_FC = 1.59, p = 6.42 x 10^-6^), a GPCR expressed in the central nervous system^63^, was also significantly upregulated in the *Mfrp^-/-^* ONL (Fig 7E), indicating changes in neural signaling pathways. Surprisingly, *Cebpd* (log_2_FC = 1.68, p = 2.43 x 10^-5^) was significantly upregulated in *Mfrp^-/-^* ONL (Fig 7E), consistent with findings in snRNA-seq analysis of rod photoreceptors (Fig 4A). Asparagine synthetase (*Asns*) (log_2_FC = 1.66, p = 0.00057), was also significantly upregulated in *Mfrp^-/-^*ONL (Fig 7E), likely indicating increased metabolic stress or nutrient deprivation^64^.

In contrast, significantly upregulated genes identified in 28d Wt ONL include *4933440N22Rik*, a long non-coding RNA that plays a role in retina development and stress responses^36^; *Vmn1r210*, a member of the vomeronasal receptor family that may contribute to immune-related signaling or cellular communication pathways^65^; *Krt14*, a cytoskeletal intermediate filament protein that may be important for maintaining photoreceptor integrity^66^; and *Serpine3,* a serine protease inhibitor, which suggests active regulation of extracellular matrix remodeling and neuroprotection^67^ (Fig 7E). The upregulation of these genes in 28d Wt mice suggests protective transcriptional processes that maintain the retinal architecture and suppresses cellular stress, which appears disrupted in *Mfrp^-/-^* ONL.

Pathway enrichment analysis of the differentially expressed genes further pinpointed specific biological pathways that were significantly perturbed in the *Mfrp^-/-^* ONL (Fig 7F). These include cholesterol biosynthesis, cellular response to nutrient starvation, and amino acid regulation of the mTORC1 pathway (Fig 7F). Mitochondrial pathways, such as respiratory electron transport, cristae formation, and formation of ATP by chemiosmotic coupling, were also marked affected (Fig 6F). Additionally, disruption of canonical pathways critical for rod photoreceptor function, including retinoid cycle and transmission across chemical synapses, was observed (Fig 7E). These findings indicate a profound impairment of both metabolic and signaling networks necessary for photoreceptor survival and function in the absence of *Mfrp*.

## DISCUSSION

MFRP-associated retinitis pigmentosa (MFRP-RP) is a clinically and genetically heterogeneous retinal dystrophy characterized by early-onset photoreceptor degeneration. Mouse models have played a critical role in elucidating the retinal defects seen in humans. Two previously characterized mouse models, rd6 and rdx, carrying spontaneous mutations in the *Mfrp* gene, exhibit phenotypes resembling human MFRP-RP, including retinal flecks and progressive thinning^12,17^. We recently described a third model harboring the patient-derived *c.498_499insC* mutation, which also exhibits retinal pathology similar to *rd6* and *rdx* mice [16]. While *Mfrp* transcript levels were elevated in *rd6* and *rdx* retinas [4], MFRP protein remains undetectable in all models^12,17^, raising questions about the impact of truncated or non-functional MFRP products. This ambiguity is mirrored in the clinical heterogeneity observed in MFRP-RP patients^5–7,10,14,68,69^, indicating the need for a clean knockout model to understand the direct consequences of complete MFRP loss.

To address this, we developed and characterized a Mfrp-/-mouse model that exhibits a fully penetrant, progressive retinal degeneration phenotype. Fundus autofluorescence imaging revealed early and increasing accumulation of autofluorescent puncta in Mfrp-/-mice beginning at 1–2 months of age, which are not seen in WT controls. These puncta are known to be reminiscent of bisretinoid accumulation, suggesting metabolic dysregulation in photoreceptors and are consistent with degeneration seen in other RP models such as rd1 and rd10^19,70^. OCT imaging and histological analysis revealed marked retinal thinning, especially of the ONL, and disruption of photoreceptor outer segments starting at P28 and culminating in near-total ONL loss by 12 months. These structural changes were accompanied by a sharp decline in ERG responses by 2 months, confirming early functional impairment. Consistent with these findings, immunohistochemistry and qRT-PCR analyses demonstrated progressive loss of opsin expression in both rods and cones. Mfrp-/-retinas showed significantly reduced Rho, Opn1mw, and Opn1sw expression at both P28 and 2.5 months, indicating a loss of both rod and cone outer segments and suggesting that MFRP is essential for photoreceptor structural maintenance.

To gain deeper insight into the molecular mechanisms driving photoreceptor degeneration in MFRP-RP, we performed single-nucleus RNA sequencing (snRNA-seq), which revealed early and widespread transcriptomic dysregulation in both rod and cone photoreceptors as early as postnatal day 28 (P28). Among the most notable changes was the altered expression of stress-response and pro-degenerative genes, including Cebpd and Egr1. EGR1, a transcription factor known to regulate inflammatory signaling and transcriptional reprogramming during retinal injury, is typically upregulated in degenerating rods and cones in other RP models such as *rd10*, where it ranks among the most strongly induced transcriptional regulators ^70^. Similar to the findings in the *rd10* model, Egr1 was upregulated specifically in cones at 2.5 months in our MFRP-RP model, suggesting a fundamentally common mechanism and possibly indicating the upregulation of protective or adaptive signaling in primary cone degeneration.

This distinct EGR1 expression pattern aligns with the broader transcriptomic shifts we observed, including downregulation of genes essential for photoreceptor structure and function such as Atp1b1, Gpm6a, Timp3, and Gnat1. These genes are involved in phototransduction, ion transport, oxidative metabolism, and extracellular matrix maintenance pathways critical for photoreceptor viability. By 2.5 months, we observed significant transcriptional dysregulation along with continued suppression of identity-maintaining and neurodevelopmental regulators like Pax6, Gpm6a, and Egr1, and upregulation of apoptosis and stress-associated genes such as Dkk3 and Unc5d. Pathway enrichment analysis revealed early activation of inflammatory and cellular stress pathways, followed by widespread suppression of synaptic, metabolic, and developmental pathways, reflecting a collapse of protective gene networks necessary for photoreceptor survival.

To determine whether these transcriptional changes were accompanied by epigenomic reprogramming, we performed sciATAC-seq, which revealed dynamic and cell type–specific alterations in chromatin accessibility in *Mfrp^-/-^* rods and cones. At P28, accessible chromatin regions were enriched for binding motifs of transcription factors associated with early stress responses and inflammation, including ELK4, FOSL2::JUND (AP-1 complex), KLF15, and STAT3. By 2.5 months, chromatin accessibility had shifted toward motifs for both photoreceptor identity regulators (CRX, OTX2) and injury-associated transcription factors (SMAD::SMAD3, FOS::JUN). This progression from stress-responsive to degeneration-associated and identity-loss signatures implicates a coordinated epigenomic shift underlying the observed transcriptional decline.

Together, these findings highlight a complex, stage-specific regulatory cascade where early stress signaling transitions into transcriptional silencing of photoreceptor maintenance programs. These molecular events are likely driven not only by cell-intrinsic degeneration but also by dysfunction in the retinal pigment epithelium (RPE), given the predominant expression of MFRP in the RPE and its role in RPE apical membrane integrity through interactions with ADIPOR1 and KCNJ13 ^71^. Although our study focused on the neural retina, the progressive degeneration of photoreceptors likely reflects both intrinsic photoreceptor dysfunction and disrupted RPE-photoreceptor interactions. These data support a dual role for MFRP in maintaining photoreceptor and RPE homeostasis through shared molecular pathways, and further suggest that targeting early transcriptional regulators such as EGR1 may provide a therapeutic window for intervention.

To better resolve the spatial context of these molecular disruptions, we applied spatial transcriptomics using GeoMx DSP, which revealed ONL-specific transcriptional alterations that complemented and validated the snRNA-seq findings. In the *Mfrp-/-* ONL, genes involved in GPCR signaling (*Gnb2, Gnb3, Gpr162*) and inflammatory responses (*Cebpd, Asns*) were upregulated, whereas protective genes such as *Krt14* and *Serpine3* were enriched in wild-type ONL. Pathway enrichment analysis further demonstrated that the *Mfrp^-/-^* ONL was significantly impaired in metabolic processes, cholesterol biosynthesis, and synaptic transmission. These results reinforce the notion that photoreceptor degeneration in MFRP-RP arises from a convergence of disrupted intracellular signaling, inflammation, and metabolic failure. These findings were not only consistent with, but also extended beyond, what was observed in the dissociated snRNA seq data.

Our transcriptomic findings in *Mfrp^-/-^* retinas share key similarities with other well-characterized RP models, including rd1 and rd10, which carry mutations in *Pde6b*^19,70^. In these models, rods degenerate rapidly, followed by secondary cone death, a pattern which is observed in Mfrp^-/-^ mice. Notably, genes such as Cebpd, Egr1, and Malat1, which were upregulated in our single-cell transcriptomic analysis of *Mfrp^-/-^*rods and cones, have also been reported as early markers of photoreceptor stress in rd10 and rd1 retinas ^19,70^. These genes are associated with inflammatory signaling, cellular stress, and transcriptional reprogramming. Additionally, downregulation of synaptic and metabolic genes such as *Atp1b1* and *Gnat1* observed in *Mfrp^-/-^*photoreceptors has been similarly reported in rd1/rd10 models, reflecting a shared molecular cascade of degeneration involving disrupted phototransduction, altered ion homeostasis, and mitochondrial dysfunction. However, while rd1/rd10 models show extremely rapid degeneration within the first month of life, *Mfrp^-/-^* mice exhibit a slightly slower, yet still early-onset progression, allowing for a broader window to study intermediate stages of degeneration. Importantly, the multi-omic integration in *Mfrp^-/-^* mice provides insights not only into transcriptional dysregulation but also into chromatin remodeling and spatial gene expression, which remain underexplored in other RP models. These comparisons position *Mfrp^-/-^* mice as a complementary and mechanistically informative model for investigating both gene-specific and convergent pathways in inherited retinal degenerations.

Taken together, our multi-omic analysis of *Mfrp^-/^*^-^ retinas reveals that MFRP is essential for maintaining photoreceptor identity, chromatin stability, and transcriptional homeostasis. Loss of Mfrp triggers an early stress response in rods, followed by cone degeneration, through distinct but overlapping transcriptomic and epigenomic programs. These insights not only validate the *Mfrp^-/-^*mouse as a robust model of MFRP-RP but also uncover shared degeneration mechanisms with other RP forms. Our study highlights new molecular targets and pathways for potential therapeutic intervention in inherited retinal diseases.

## METHODS

### Animal models

Homozygous *Mfrp^-/-^* mice (C57BL/6 background) were generated by breeding with previously established homozygous *Mfrp^KI/KI^* mice^16^ with a ISL-1 Cre/Tyrosinase related protein 1 (Trp1) gene promoter to target the DNA region between exons 3-9 of the *Mfrp* gene flanked by LoxP regions. The resulting *Mfrp^-/-^* mice with biallelic knockout was confirmed by genotyping using gene-specific primers for PCR (Supplementary Fig 1C). C57BL/6 background Wt mice were used as controls for studies. Wt and *Mfrp^-/-^* mice were maintained under standard conditions with a 12-hourlight/12-hour dark cycle with access to food and water. Both male and female mice were used in this study. All animal procedures were approved by the Institutional Animal Care and Use Committee (IACUC)

### Fundus and OCT

*In vivo* imaging was performed on mice anesthetized with intraperitoneal injections of ketamine (93 mg/kg) and xylazine (8mg/kg). Pupils were treated with 0.5% Proparacaine hydrochloride ophthalmic solution USP, 0.5% followed by dilation with 10%% phenylephrine hydrochloride ophthalmic solution and 1% tropicamide ophthalmic solution. Spectralis™ HRA + OCT (Heidelberg Engineering, Germany) was used to perform fundus autofluorescence imaging and spectral domain optical coherence tomography (SD-OCT) of Wt and *Mfrp^-/-^* mice.

### Western blotting

Mouse eyes were enucleated and the RPE-choroid were physically separated from the neural retina. Pooled RPE-choroid were lysed in RIPA buffer containing protease and phosphatase inhibitor cocktail (Millipore, Sigma, Cat# 11697498001) and incubated on ice for 20 min with intermittent vortexing. Lysates were prepared by centrifugation at 14,000rpm for 20 min at 4°C to remove the cell debris. The supernatant was collected, and protein concentration was determined using a Bradford assay. Equal amounts of protein were loaded onto a Novex™ WedgeWell™ 4 to 20%, Tris-Glycine gels (ThermoScientific, cat# XP04200) and transferred onto a PVDF membrane (ThermoScientific, cat#: 88518). Membranes were blocked in 5% milk for 1hr then probed with mouse anti-MFRP (1:1000; R&D systems, cat#AF3445) and mouse anti-β-actin (1:5000; Sigma, cat# A2228) primary antibodies in 3% milk overnight. Secondary antibodies used were donkey anti-mouse IgG (1:3,000; ThermoScientific, cat# 31430) and donkey anti-rabbit IgG (1:3,000; ThermoScientific, cat# A16035). β-actin which was used as a loading control. Rabbit polyclonal antibodies against phosphorylated and total AMPK, ACC (cell signaling cat#. 2535, 3661 respectively) were also used.

### Histology and Immunohistochemistry

6-8µm Cryosections of Wt and *Mfrp^-/-^*retinas were used to perform histology and immunohistochemistry as described previously^72^. The following primary antibodies were used for staining: Opsin (1:200, Millipore Sigma, cat# AB5405), OPN1SW (1:200, Santa Cruz Biotechnology, cat# sc-14363) and Rhodopsin (1:200, Abcam, cat# ab3424). Secondary antibodies include donkey anti-goat AlexaFluor 488 (Invitrogen, cat# A11055) and donkey anti-rabbit AlexaFluor 555 (Invitrogen, cat# A31572). Histology images were captured using an EVOS FL Auto Imaging System (Life Technologies, cat# AMAFD1000) and immunofluorescence mages were captured using a ZEISS LSM 800 confocal microscope.

### Single-nucleus ATAC-seq data generation

Combinatorial barcoding single-nucleus ATAC-seq was provided by the Center for Epigenomics, UC San Diego and performed as previously described ^73–75^. Briefly each frozen mouse retina was homogenized using a glass dounce homogenizer in 1 mL of douncing buffer consisting of 0.25 M sucrose (S1888, Sigma), 25 mM KCl (AM9610G, Invitrogen), 5 mM MgCl2 (194698, Mp Biomedicals Inc.), 10 mM Tris-HCl pH 7.5 (15567027, Thermo Fischer Scientific), 1 mM DTT (D9779, Sigma), protease inhibitor (05056489001, Roche), 0.1% Triton X-100 (T8787-100ML, Sigma), and 0.2 U/µL RNasin RNase inhibitor (PAN21110, Promega) in molecular biology grade water (46000-CM, Corning). Nuclei were spun at 300g x 5 minutes and resuspending in 1 ml nuclei permeabilization buffer (10 mM Tris-HCL [pH 7.5], 10 mM NaCl, 3 mM MgCl2, 0.1% Tween-20 [Sigma], 0.1% IGEPAL-CA630 [Sigma] and 0.01% Digitonin [Promega] in water) Nuclei suspension was incubated for 10 min at 4°C and filtered with 30 μm filter (CellTrics). Nuclei were pelleted with a swinging bucket centrifuge (500 x g, 5 min, 4°C; 5920R, Eppendorf), resuspended in 500 µL high salt tagmentation buffer (36.3 mM Tris-acetate (pH = 7.8), 72.6 mM potassium-acetate, 11 mM Mg-acetate, 17.6% DMF) and counted using a hemocytometer. Concentration was adjusted to 2000 nuclei/9 µL, and 2000 nuclei were dispensed into each well of one 96-well plate. For tagmentation, 1 µL barcoded Tn5 transposomes ^76^ was added using a BenchSmart 96 (Mettler Toledo), mixed five times and incubated for 60 min at 37°C with shaking (500 rpm). To inhibit the Tn5 reaction, 10 µL of 40 mM EDTA were added to each well with a BenchSmart 96 (Mettler Toledo) and the plate was incubated at 37°C for 15 min with shaking (500 rpm). Next, 20 µL 2 x sort buffer (2% BSA, 2 mM EDTA in PBS) was added using a BenchSmart 96 (Mettler Toledo). All wells were combined into a FACS tube and stained with 3 µM Draq7 (Cell Signaling). Using a SH800 (Sony), 20 2 n nuclei were sorted per well into eight 96-well plates (total of 768 wells) containing 10.5 µL EB (25 pmol) primer i7, 25 pmol primer i5, 200 ng BSA (Sigma). Preparation of sort plates and all downstream pipetting steps were performed on a Biomek i7 Automated Workstation (Beckman Coulter). After addition of 1 µL 0.2% SDS, samples were incubated at 55°C for 7 min with shaking (500 rpm). 1 µL 12.5% Triton-X was added to each well to quench the SDS. Next, 12.5 µL NEBNext High-Fidelity 2 × PCR Master Mix (NEB) were added and samples were PCR-amplified (72°C 5 min, 98°C 30 s, (98°C 10 s, 63°C 30 s, 72°C 60 s)×12 cycles, held at 12°C). After PCR, all wells were combined. Libraries were purified according to the MinElute PCR Purification Kit manual (Qiagen) using a vacuum manifold (QIAvac 24 plus, Qiagen) and size selection was performed with SPRI Beads (Beckmann Coulter, 0.55x and 1.5x). Libraries were purified one more time with SPRI Beads (Beckmann Coulter, 1.5x). Libraries were quantified using a Qubit fluorimeter (Life technologies) and the nucleosomal pattern was verified using a Tapestation (High Sensitivity D1000, Agilent). The library was sequenced on a NovaSeq6000 or NextSeq500 sequencer (Illumina) using custom sequencing primers with following read lengths: 50 + 10 + 12 + 50 (Read1 + Index1 + Index2 + Read2).

### Single-nucleus RNA-seq data generation

Single nucleus RNA-seq was provided by the Center for Epigenomics, UC San Diego using the Droplet-based Chromium Single-Cell 3’ solution (10x Genomics, v3 chemistry) ^77^. Briefly, nuclei isolated as described in single nucleus ATAC-seq methods were suspended in 400 µL of sort buffer (1 mM EDTA 0.2 U/µL RNase inhibitor (Promega, N211B), 2% BSA (Sigma-Aldrich, SRE0036) in PBS) and stained with DRAQ7 (1:100; Cell Signaling, 7406). 75,000 nuclei were sorted using a SH800 sorter (Sony) into 50 µL of collection buffer consisting of 1 U/µL RNase inhibitor in 5% BSA; the FACS gating strategy sorted based on particle size and DRAQ7 fluorescence. Sorted nuclei were then centrifuged at 1000 rcf for 15 min (4°C, run speed 3/3) and supernatant was removed. Nuclei were resuspended in 35 µL of reaction buffer (0.2 U/µL RNase inhibitor (Promega, N211B), 2% BSA (Sigma-Aldrich, SRE0036) in PBS) and counted on a hemocytometer. 12,000 nuclei were loaded onto a Chromium Controller (10x Genomics). Libraries were generated using the Chromium Single-Cell 3′ Library Construction Kit v3 (10x Genomics, 1000075) with the Chromium Single-Cell B Chip Kit (10x Genomics, 1000153) and the Chromium i7 Multiplex Kit for sample indexing (10x Genomics, 120262) according to manufacturer specifications. CDNA was amplified for 12 PCR cycles. SPRISelect reagent (Beckman Coulter, B23319) was used for size selection and clean-up steps. Final library concentration was assessed by Qubit dsDNA HS Assay Kit (Thermo-Fischer Scientific) and fragment size was checked using Tapestation High Sensitivity D1000 (Agilent) to ensure that fragment sizes were distributed normally about 500 bp. Libraries were sequenced using the NextSeq500 and a NovaSeq6000 (Illumina) with these read lengths: 28 + 8 + 91 (Read1 + Index1 + Read2).

### snRNA-seq Data Processing and Analysis

Data was processed and mapped to mm10 using Cell Ranger (10x Genomics). Raw count matrices were processed to remove ambient and background RNA using CellBender. Potential doublets were identified and removed with DoubletFinder. The resulting nine datasets were merged with Seurat. Quality control (QC) filtering retained nuclei with 200 < *nFeature_RNA* < 6,000 and mitochondrial transcript content (*percent.mt*) < 2%. Data were normalized using SCTransform, followed by principal component analysis (PCA) and batch correction with Harmony (*RunHarmony()*). Clustering was performed in Seurat using *FindNeighbors()*, *FindClusters()*, and *RunUMAP()* with 50 principal components and a resolution parameter of 0.6. After removing doublets and low-quality cells, 57,420 high-quality nuclei were retained for downstream analysis. Clusters were annotated based on canonical marker gene expression, yielding 19 distinct retinal cell types. Differential gene expression analysis was conducted using DESeq2, with differentially expressed genes (DEGs) defined as those having an adjusted *p* < 0.05 and absolute fold-change > 1.5. Functional enrichment analysis was performed using clusterProfiler, with DEGs as input to identify significantly enriched pathways.

### scATAC-seq Data Processing and Analysis

sciATAC-seq was processed and mapped to mm10 as previously described ^78^. Nine individual datasets were merged into a single Seurat object. Doublets were identified and removed using Scrublet with an expected doublet rate of 6%. Quality control (QC) thresholds included nucleosome signal > 4, transcription start site (TSS) enrichment > 5, and > 1,000 unique usable fragments per nucleus. The sciATAC-seq data were processed using Signac and Seurat, including TF-IDF normalization, batch correction with Harmony, and clustering (35 dimensions, resolution = 1.5). A total of 71,335 high-quality nuclei passed QC. Cluster annotation was performed using Seurat’s label transfer workflow, applying *FindTransferAnchors()* and *TransferData()* to project cell-type labels from the matched snRNA-seq dataset onto the sciATAC-seq data, resulting in 16 annotated retinal cell types. Differential Transcription factor biding MOTIF analysis was performed using ChromVar and presto, applying the *wilcoxauc.Seurat()* function to run Wilcoxon rank-sum testing and compute area under the ROC curve (auROC) scores on chromVAR motif deviation values.

### Spatial transcriptomic data generation and analysis

Whole-transcriptome spatial profiling was performed using the GeoMx DSP platform (NanoString Technologies). Retinal sections were incubated with the Mouse Whole Transcriptome Atlas probes alongside morphology markers (SYTO13 for nuclei and antibody-based markers for region identification). Regions of interest (ROIs) were selected manually based on retinal layer architecture, focusing specifically on the outer nuclear layer (ONL). UV photocleavage of oligonucleotide tags was performed for each ROI, followed by oligo collection and library preparation according to the manufacturer’s protocol. Oligonucleotide libraries were prepared using the GeoMx DSP Library Prep Kit and barcoded for multiplexed sequencing. Libraries were quantified by Qubit fluorometric assay and quality-checked using a Bioanalyzer. Sequencing was performed on an Illumina NextSeq or equivalent platform, aiming for a minimum depth of 1 million reads per ROI. Raw sequencing reads were demultiplexed, aligned, and quantified using the GeoMx NGS pipeline. Normalization was performed using the Q3 method. Differential expression analysis between 28d *Mfrp^-/-^* and Wt ONL ROIs was conducted using the NanoString GeoMx DSP analysis suite. For differential gene expression, statistical significance was determined using a Benjamini-Hochberg false discovery rate (FDR) correction with an adjusted p-value threshold of 0.05 unless otherwise indicated.

### Statistical analysis

To determine statistical significance, GraphPad Prism software was used to perform a one-sample *t-*test for the comparison of Wt vs. *Mfrp^-/-^* mice. Significance was set at \**p* < 0.05, \*\**p* < 0.01, \*\*\**p* < 0.001 and \*\*\*\**p*<0.0001. ns = not significant = *p* > 0.05.

