## Supplementary figures and images for "Multi-omic analysis of photoreceptor alterations during early-onset retinal degeneration in *Mfrp^-/-^* mice"

### Supplementary figure 1

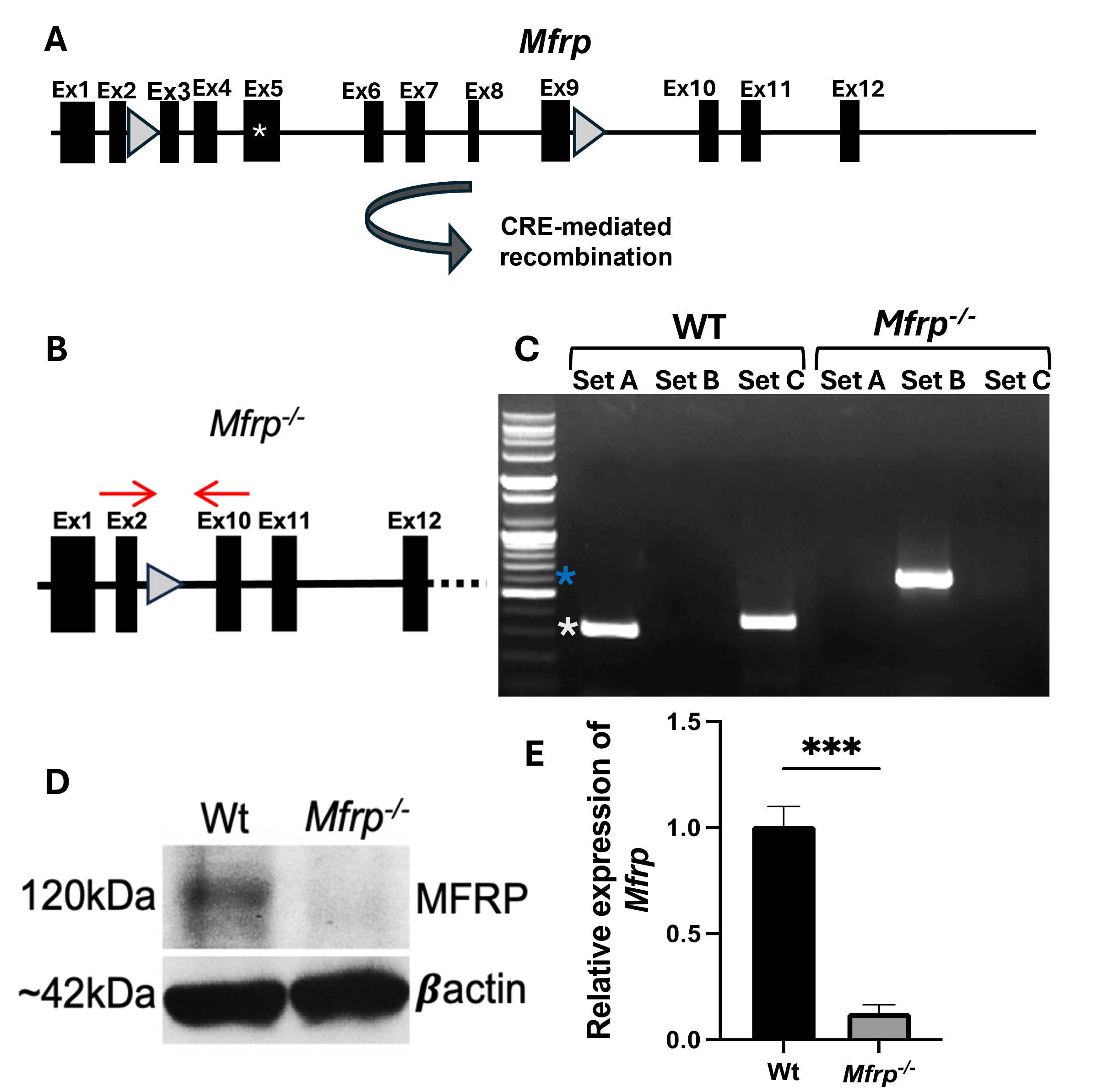
